# Balanced Permeability Index is a Strong Predictor of Intestinal Absorption and Oral Bioavailability for Heterobifunctional Ligand-Directed Degraders

**DOI:** 10.1101/2025.03.19.644178

**Authors:** Javier L. Baylon, Matthew J. Chalkley, Tony Siu, Wilson Shou, Yongnian Sun, Xianmei Cai, Anthony Paiva, Shivani Patel, Tatyana Zvyaga, Dahlia R. Weiss

**Affiliations:** Bristol-Myers Squibb Company, San Diego, CA 92121; Bristol-Myers Squibb Company, Redwood City, CA 94063; Bristol-Myers Squibb Company, Cambridge, MA 02141; Bristol-Myers Squibb Company, Lawrence Township, NJ 06848

**Keywords:** Heterobifunctional degrader, Proteolysis targeting chimeras, LDD, BPI, Bioavailability

## Abstract

Ligand-directed degraders (LDDs) are heterobifunctional molecules that degrade proteins by engaging the ubiquitin-protein ligase (E3) system. LDDs consist of a target-engaging moiety, an E3 ligase-binding moiety and a bridging linker. Due to their size and physicochemical complexity these molecules do not adhere to well-established rules of lead optimization. The optimization of passive permeability remains a key challenge to develop orally bioavailable LDDs. To overcome this challenge, in this study we demonstrate that the Balanced Permeability Index (BPI)—a new metric that combines size, polarity and lipophilicity—is highly predictive of oral bioavailability for LDDs. Here, we introduce an additional parameter—called smallest maximum intramolecular distance (SMID)—to the original BPI index to account for cross sectional area of LDDs, termed BPI_LDD_. With this new parameter, BPI_LDD_ can differentiate oral bioavailability of LDDs in our dataset more effectively than polarity, lipophilicity, or size separately. In addition, BPI_LDD_ is also more effective at identifying orally bioavailable LDDs than some *in vitro* measurements of cell permeability that traditionally inform bioavailability. This finding opens the possibility of employing BPI_LDD_ for the design and optimization of orally bioavailable LDDs to improve their drug metabolism and pharmacokinetics properties.

---

Targeted protein degradation (TPD) has emerged as an exciting new modality that seeks to eliminate disease causing proteins, especially in cases where those targets are considered traditionally ‘undrugabble’ or have critical scaffolding functions. Heterobifunctional degraders or Ligand-directed degraders (LDDs) were introduced over 20 years ago and have since matured into a viable clinical modality.^3-10^ LDDs consist of three main segments: a target-binding moiety, a linker and an E3 ligase binding moiety.^3, 11^ Since their initial inception, LDDs have demonstrated great therapeutic potential to degrade challenging targets, sustained efficacy through an event driven pharmacology rather than the traditional occupancy driven pharmacology.^12-14^ However, due to their molecular complexity, rational design of LDDs remains a challenging task.^15^ Although the chemical space for ligands targeting E3 ligases is limited to only a few molecules,^16^ the combinatorial nature of assembling the entire LDD makes the overall chemical space large.^17, 18^ Moreover, the chemical complexity of LDDs limits the number of molecules that can be produced during a typical design-make-test cycle. These features render LDDs unsuitable for computational approaches such as machine learning (ML) for design and optimization due to the sparse datasets and the lack of transferability of ML results across diverse LDD chemical spaces.^19^

Compounding this challenge, LDDs have physicochemical properties that are significantly different from traditional small molecules.^20^ LDDs present high molecular weight (MW), large polar surface area (PSA) and many rotatable bonds. These “beyond rule of 5” (bRo5) properties have a direct and often negative impact on the drug metabolism and pharmacokinetics (DMPK) properties of LDDs.^21^ Moreover, LDDs are known to behave poorly in a number of *in vitro* assays typically applied in early stages of development of Ro5 molecules obscuring efforts to predict oral bioavailability.^22, 23^

Thus, several strategies have been proposed to predict and optimize oral bioavailability of bRo5 compounds using key physicochemical descriptors, such as molecular weight (MW), lipophilicity (ClogP), hydrogen bond donors (HBD) and acceptors (HBA) and PSA,^24-26^ as well as composites scores combining some of those properties.^27^ More recently, chromatographic approaches have shown promise to aid development of permeable LDDs including exposed polar surface area (EPSA),^28^ ChameLogK,^29^ and ChameLogD.^29^ Despite these advancements in the design of bRo5 molecules, new strategies are needed to rationally understand and optimize the physiochemical properties of LDDs to achieve orally bioavailable compounds and unlock the potential of this emerging therapeutic modality.

In this study we show that the recently introduced Balanced Permeability Index (BPI)^30^ is well correlated to favorable oral absorption and bioavailability for a series of LDDs. BPI was originally introduced to optimize *in vitro* permeability and efflux of molecules using by combining size, polarity, and lipophilicity —the key properties driving passive permeability and P-gp efflux—into a single metric.^30^ BPI is defined as:

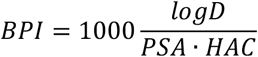

where LogD is a measure of lipophilicity, PSA is the polar surface area and HAC is the heavy atom count of the molecule. For the experimental BPI, these values are measured using two chromatographic assays, exposed polar surface area (EPSA) and LogD at pH 7.4 (LogD_7.4_) and normalized by HAC.

To investigate whether BPI can serve as a predictor of LDD oral absorption, we focused on the F_a*_F_g_. F_a*_F_g_ refers to the fraction of a drug absorbed in the intestine that reaches systemic circulation, defined as^31-33^:

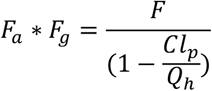

where F is oral bioavailability, Cl_p_ is total plasma clearance and Q_h_ is hepatic blood flow. To this end we selected a series of compounds consisting of 129 cereblon-recruiting LDDs from different internal programs for which experimentally measured LogD at pH 7.4 and experimentally determined exposed polar surface area (EPSA) data was available. The details of these assays have been reported elsewhere.^34, 35^ To ensure consistency, F_a*_F_g_ was calculated (Mouse, Qh = 90 mL/min/kg) from compounds where mouse oral bioavailability is available. Standard formulation and dosing were not enforced for the F_a*_F_g_ data set to avoid significantly reducing the size of the datasets. Instead, formulation and dosing were chosen independently in each study by subject matter experts. Based on previous research that set a threshold of F_a*_F_g_ > 0.25 for desirable absorption, the compounds were categorized as poor and high F_a*_F_g_. ^2^ Out of the total compounds, 94 fell into the poor absorption category, while 35 fell into the high absorption category.

The ROC plot in Figure 1 demonstrates that BPI can enrich compounds with high F_a*_F_g_ (defined as F_a*_F_g_>25%) and outperforms its individual components, albeit with very modest improvement over EPSA (AUC of 0.7 vs 0.67 respectively).

**Figure 1.**
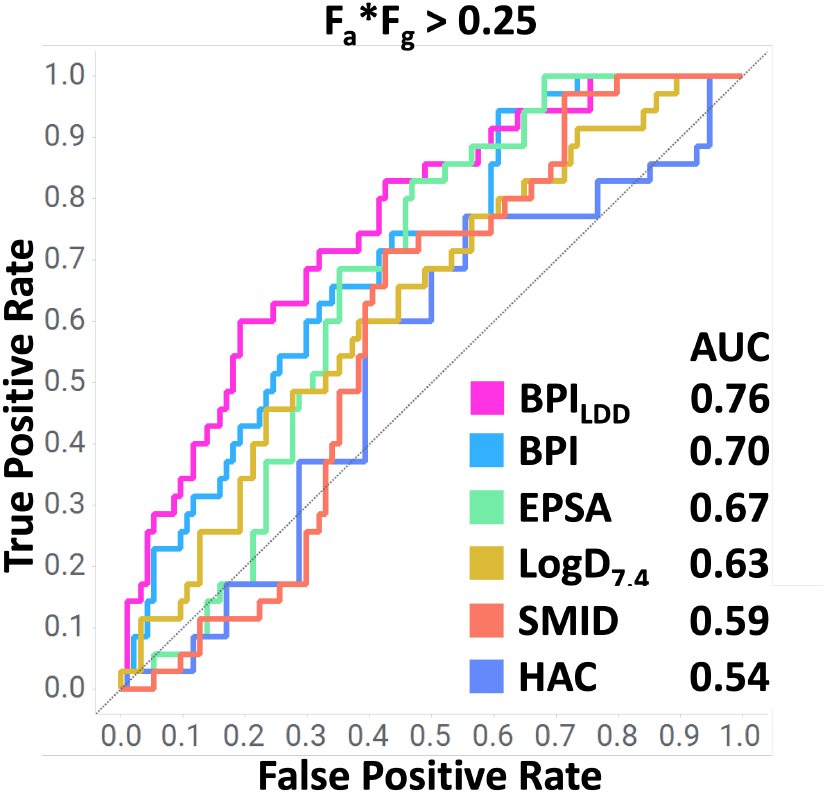
Comparison of BPI and other parameters to differentiate LDDs with desirable intestinal absorption F_a*_F_g_ > 0.25. Area under curve (AUC) for each parameter is shown in the legend.

The relationship between a molecule’s shape and its passive membrane permeability is complex and multifaceted, however, molecules that can adopt a more rigid and rod-like conformation tend to permeate more easily, as they displace fewer membrane lipids by presenting a small cross-sectional area during passage. This aspect may be particularly relevant for LDD molecules, which have the potential to adopt rod-like conformations at the cost of exposing polar residues, which would favor a more spherical conformation to bury these polar groups. We hypothesized that LDD molecules capable of satisfying both requirements, minimizing surface polarity while maintaining a rod-like conformation, would exhibit enhanced absorption. To quantify this balance, we introduce a modified version of the BPI metric for LDDs that considers the length of the molecule, defined as:

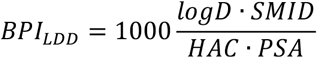

where SMID is the smallest maximum intramolecular distance, an *in silico* parameter used to characterize the longest-axis of the smallest conformation that a molecule can adopt.^36^ The SMID was calculated using a publicly available workflow that has been previously described in detail elsewhere.^36^ By including SMID in the numerator, the contribution of PSA in the denominator effectively becomes a cross-sectional “polar” area, which has been shown to be highly predictive of permeability.^34, 35^ While this relationship was driven empirically, we hypothesized that this correction may improve performance compared to Ro5 small molecules. Indeed, the new metric BPI_LDD_ provides significant enrichment relative to EPSA (AUC 0.76 vs 0.67). In addition, both BPI metrics offer early enrichment (higher rate of true positives at a low rate of false positives in the first 20% of the dataset) compared to the other parameters (Figure 1). This ROC analysis demonstrates the advantage of using BPI_LDD_ to identify LDDs with high F_a*_F_g_ in our dataset.

Next, we examined the distributions of BPI, BPI_LDD_, and their components across poorly and highly absorbed compounds in our dataset (Figure 2 and Figure S1). The analysis revealed that BPI_LDD_ is a better classifier of F_a*_F_g_ than each of its components individually. Importantly, statistical analysis indicated that all these relationships except HAC and SMID are significant at differentiating between the different ranges of oral absorption (Kruskal-Wallis one-way ANOVA test, p<0.05, full results available in Supporting Information).

**Figure 2.**
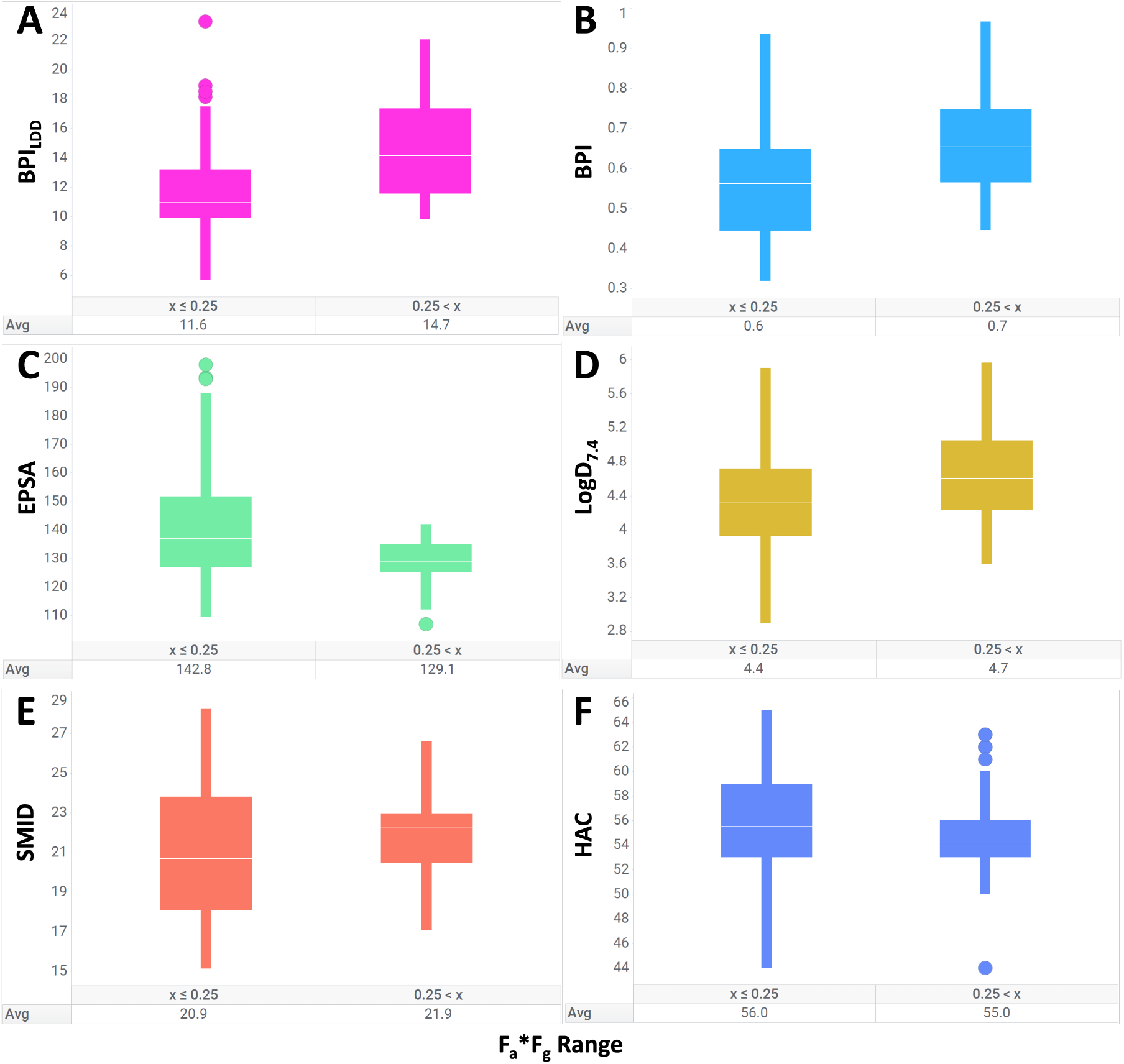
BPI_LDD_ is a strong predictor of oral bioavailability. Distribution of (A) BPI_LDD_, (B) BPI and its individual components (C) EPSA, (D) LogD*7*.*4*, (E) SMID, and (E) heavy atom count (HAC) for the LDD dataset against oral bioavailability. Table under each plot indicates the average value of the parameter for the compounds in each range.

High F_a*_F_g_ in our dataset is correlated with low EPSA and smaller number of heavy atoms as expected. However, the operating ranges of these parameters are not easily discernible. For example, the average HAC for poor and high F_a*_F_g_ in our dataset only varies by one atom (56 versus 55). In a lead optimization campaign for LDDs, it is common to have small operating ranges in terms of size, polarity, and lipophilicity. This is because the ligase-binding and target-binding moieties typically remain relatively unchanged, and linker lengths also fall within a narrow range. In contrast, by a including a term that considers SMID into the BPI_LDD_, the trend is clearer (Figure 2A) and the dynamic range is larger, with an average BPI_LDD_ of 11.6 and 14.7 for the low and high F_a*_F_g_ bins respectively. These findings suggest that permeable, highly absorbed LDDs successfully balance the trade-off between minimizing polarity and attaining a more elongated, rod-like shape, even within congeneric series of LDD with similar chemical structures. This in turn presents a greater dynamic range for optimization within a chemical series.^30^ Based on the data herein, which originated from four distinct cereblon-targeting LDD programs (Figure S2), an emerging guideline for LDDs is to strive for BPI_LDD_ > 14 to achieve moderate to high oral bioavailability. However, it should be noted that simply having a BPI_LDD_ value above 14 does not guarantee desirable F_a*_F_g_, as oral absorption and bioavailability are influenced by various other factors such as stability, dissolution rates in the gut, interactions with transporter proteins, and non-specific protein binding, among others. Additionally, it is important to consider that the specific cutoff for high BPI_LDD_ may vary depending on the chemotype of a particular LDD series.^19^ Therefore, it is strongly recommended to analyze the LDD series of interest to derive a specific cutoff for high BPI_LDD_ to reliably predict high oral bioavailability.

To investigate whether the experimentally derived BPI_LDD_ provided additional insights relative to previously proposed computed physiochemical predictors of LDD bioavailability, we performed ROC analysis for metrics that have been to correlate with LDD permeability or oral bioavailability F (Figure 3).^26^ These properties include heavy atom count (shown in Figure 2F), as well as total number of hydrogen bonds (HB, sum of HBA and HBD), MW and number of rotatable bonds (NRB). We also calculated the AbbVie multiparametric scoring function (AB-MPS) derived from the AbbVie compound collection for bRo5 compounds, which has been widely employed.^27^ The AB-MPS score is defined as:

**Figure 3.**
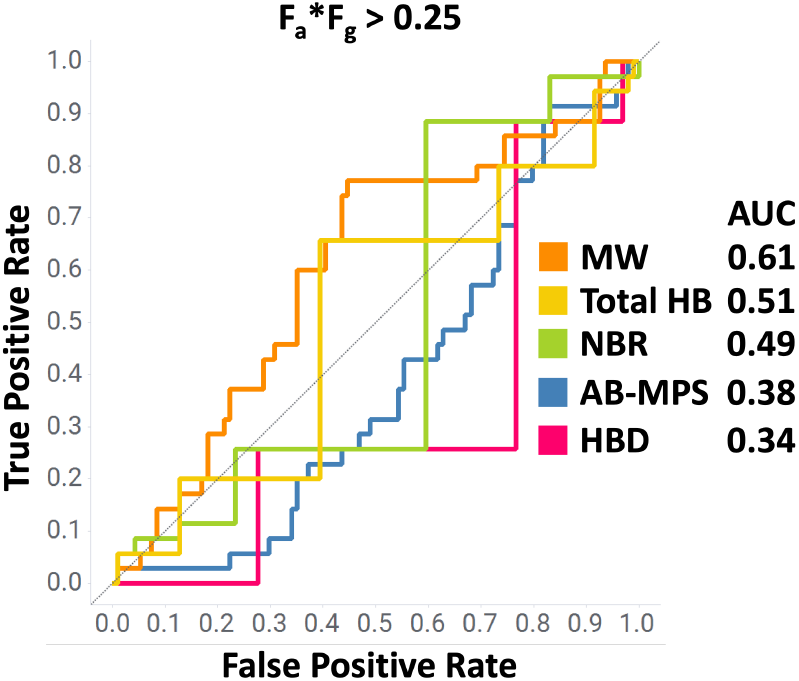
ROC analysis shows that physicochemical properties associated with oral bioavailability are not predictive of F_a*_F_g_ for LDDs in the dataset. None of the parameters considered, including AB-MPS, can distinguish LDDs with high F_a*_F_g_ > 0.25. AUC for each parameter is shown in the legend.

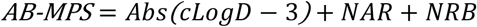

where cLogD is calculated LogD for lipophilicity, NAR is the number of aromatic rings, and NRB is the number of rotatable bonds. Notably, none of these computed molecular properties achieved the same level of classification of LDDs by F_a*_F_g_ in ROC analysis as BPI_LDD_, as evidenced by their AUC values being lower than the BPI_LDD_ AUC of 0.76. In fact, our analysis revealed that these physicochemical properties were unable to effectively identify compounds with desirable F_a*_F_g_ in our dataset.

We also investigated whether passive permeability (Papp A-to-B) in Cancer coli (Caco-2) cells is predictive of oral bioavailability for the LDDs in our set (Figure S3). It has been noted previously that LDD molecules present unique challenges in the performance of routine *in vitro* assays. For example, in our F_a*_F_g_ dataset, we found 78 molecules that had been assayed in the Caco-2 cell line. However, the mass balance recovery for these compounds was predominantly poor, with over two-thirds of the compounds (65 out of 78 compounds) failing to achieve a recovery rate greater than 80%. This indicates that for most of the compounds assayed, at least 20% of the material was unrecovered at the end of the experiment due to non-specific depletion such as metabolism in the membrane, binding to the membrane and assay materials, and so on. Out of our LDD dataset, only 13 compounds with measured F_a*_F_g_ had acceptable recovery rates from the Caco-2 assay (>80%). The Caco-2 Papp A-to-B in our 13-compound dataset was not able to differentiate the two ranges of F_a*_F_g_ defined previously (Kruskal-Wallis one-way ANOVA test, p=0.07). In fact, only one out of the 13 compounds showed Caco2 Papp A-to-B > 10 × 10^−6^ cm/sec that is typically indicative of highly permeable molecules (regardless of whether those compounds were actually orally absorbed).^30^ More importantly, the low recovery across the board severely limited the applicability of this assay to select for compounds to progress to *in vivo* PK experiments. This highlights a general limitation of permeability assays not optimized for LDDs, and emphasizes the added advantage of using BPI_LDD_, which relies solely on well-behaved chromatographic measurements of EPSA and logD.

Finally, we investigated the applicability of our new BPI_LDD_ to predict oral bioavailability (%F) for a public dataset consisting of 55 LDDs (Figure S4 and Figure S5).^1^ This dataset did not report measured LogD_7.4_ or EPSA values so we used the calculated cLogP and TPSA values provided with the dataset to predict %F (F_a*_F_g_ was not reported). Additionally, we calculated the SMID values for each of the 55 LDDs in the dataset using the pre-established workflow publicly available.^36^ Therefore, for this analysis we only used calculated and computationally derived parameters to calculate BPI_LDD_ from public sources. To perform the analysis, we defined two ranges for %F with a threshold at %F > 20 to be consistent with previously reported analysis of LDD molecules,^2^ resulting in a split of 26 and 29 LDDs with high and low %F, respectively. Unsurprisingly, the performance of BPI_LDD_ is degraded by the lack of experimental data (AUC = 0.73), likely due to underperforming contribution from cLogP. It is also unsurprising that HAC and TPSA perform similarly to the BPI metrics for enriching molecules with high oral bioavailability, and this is because this data set is comprised of multiple chemical series, including VHL and CRBN targeting LDD, with a large range of sizes and polarities. This is less reflective of a true optimization campaign which would generally compare LDD in a chemically related series. Interestingly, SMID and the cross sectional “polar” area defined as SMID/TPSA demonstrated better predictive performance at differentiating LDDs with high %F for the public dataset. This demonstrates the potential utility of including SMID/TPSA as part of a prospective tool for the development of new LDDs where experimental EPSA and LogD_7.4_ data is lacking.

In conclusion, our study demonstrates the applicability of BPI in predicting the absorption and oral bioavailability of LDDs. We observed a significant improvement in the ability to differentiate orally absorbed LDDs by adding an additional parameter to the index termed BPI_LDD_, that considered a crosssectional polar surface area correction. We used experimental LogD_7.4_ and EPSA values for our internal LDD set and calculated values for a public LDD set, showing our metrics are useful for both measured and calculated values. Additionally, this study showed that BPI_LDD_ can be used for *in vivo* predictions to fill in gaps of *in vitro* assays that are not optimized for bRo5 molecules. By considering factors such as size, lipophilicity, and cross-sectional “polar” area, the BPI_LDD_ can provide valuable guidance in the optimization of orally bioavailable LDDs.

## ASSOCIATED CONTENT

### Supporting Information

The Supporting Information is available free of charge on the ACS Publications website.

PDF file with additional plots and analysis (PDF).

Excel file with all the data points used for analyses and Kruskal-Wallis results (XLSX)

## AUTHOR INFORMATION

### Author Contributions

The manuscript was written through contributions of all authors. All authors have given approval to the final version of the manuscript.

## ACKNOWLEDGMENT

We would like to acknowledge the work of the Lead Discovery and Optimization ADME team for permeability measurements, and the Pharmaceutical Candidate Optimization team.

## ABBREVIATIONS

LDD: Ligand-directed degraders
BPI: Balanced Permeability Index
PAMPA: Parallel artificial membrane permeability assay
MDCK: Madin-Darby canine kidney
Caco-2: Cancer coli
DMPK: drug metabolism and pharmacokinetics
MW: molecular weight
ML: machine learning
HBA: hydrogen bond acceptor
HBD: hydrogen bond donor
NRB: number of rotatable bond
NAR: number of aromatic rings
BMS: Bristol-Myers Squibb Company
HAC: heavy atom count
TPSA: Topological polar surface area
EPSA: exposed polar surface area
SMID: smallest maximum intramolecular distance
Caco-2: Cancer coli
AB-MPS: AbbVie multiparametric scoring function.

---

For Table of Contents Only

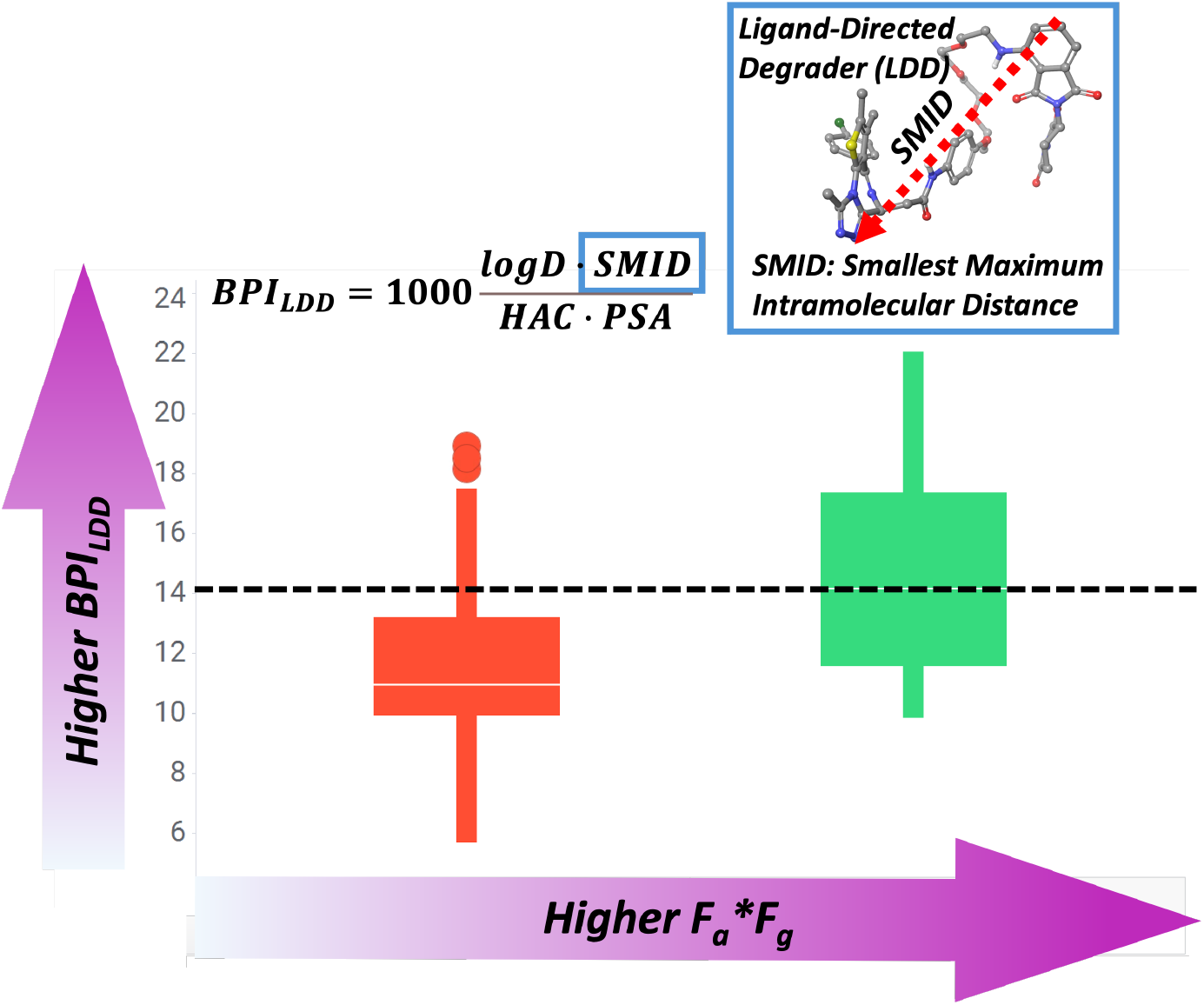

## REFERENCES

(1) Apprato, G.; Poongavanam, V.; Garcia Jimenez, D.; Atilaw, Y.; Erdelyi, M.; Ermondi, G.; Caron, G.; Kihlberg, J. Exploring the chemical space of orally bioavailable PROTACs. Drug Discov Today 2024, 103917. DOI: 10.1016/j.drudis.2024.103917 From NLM Publisher.

(2) Hornberger, K. R.; Araujo, E. M. V. Physicochemical Property Determinants of Oral Absorption for PROTAC Protein Degraders. J Med Chem 2023, 66 (12), 8281–8287. DOI: 10.1021/acs.jmedchem.3c00740 From NLM Medline.

(3) Sakamoto, K. M.; Kim, K. B.; Kumagai, A.; Mercurio, F.; Crews, C. M.; Deshaies, R. J. Protacs: chimeric molecules that target proteins to the Skp1-Cullin-F box complex for ubiquitination and degradation. Proc Natl Acad Sci U S A 2001, 98 (15), 8554–8559. DOI: 10.1073/pnas.141230798 From NLM Medline.

(4) Rathkopf, D. E.; Patel, M. R.; Choudhury, A. D.; Rasco, D. W.; Lakhani, N. J.; Hawley, J. E.; Aparicio, A.; Narayan, V.; Srinivas, S.; Runcie, K. First-in-human phase 1 study of CC-94676, a first-in-class androgen receptor (AR) ligand-directed degrader (LDD), in patients (pts) with metastatic castration-resistant prostate cancer (mCRPC). American Society of Clinical Oncology: 2024.

(5) Linton, K.; Doorduijn, J.; El-Sharkawi, D.; Mous, R.; Forconi, F.; Lewis, D.; Gleeson, M.; Riches, J.; Mckay, P.; Stevens, W. PB2331: An ongoing first-in-human Phase 1 Trial of NX-5948, an oral Bruton”s tyrosine kinase (BTK) degrader, in patients with relapsed/refractory B cell malignancies. Hemasphere 2023, 7 (S3), e29005fd.

(6) Smith, S. D.; Starodub, A.; Stevens, D. A.; Shastri, A.; Porcu, P.; Feldman, T.; Ewesuedo, R.; DeSavi, C.; Dey, J.; Agarwal, S. A Phase Study of KT-333, a Targeted Protein Degrader of STAT3, in Patients with Relapsed or Refractory Lymphomas, Large Granular Lymphocytic Leukemia, and Solid Tumors. Blood 2022, 140 (Supplement 1), 12024–12025.

(7) Mato, A.; Danilov, A. V.; Patel, M. R.; Tees, M. T.; Flinn, I. W.; Ai, W. Z.; Patel, K.; Wang, M.; O”Brien, S. M.; Nandakumar, S. A first-in-human phase 1 trial of NX-2127, a first-in-class oral BTK degrader with IMiD-like activity, in patients with relapsed and refractory B-cell malignancies. American Society of Clinical Oncology: 2022.

(8) Hamilton, E. P.; Schott, A. F.; Nanda, R.; Lu, H.; Keung, C. F.; Gedrich, R.; Parameswaran, J.; Han, H. S.; Hurvitz, S. A. ARV-471, an estrogen receptor (ER) PROTAC degrader, combined with palbociclib in advanced ER+/human epidermal growth factor receptor 2–negative (HER2-) breast cancer: Phase 1b cohort (part C) of a phase 1/2 study. J. Clin. Oncol 2022, 40, 16.

(9) Gao, X.; Burris III, H. A.; Vuky, J.; Dreicer, R.; Sartor, A. O.; Sternberg, C. N.; Percent, I. J.; Hussain, M. H.; Rezazadeh Kalebasty, A.; Shen, J. Phase 1/2 study of ARV-110, an androgen receptor (AR) PROTAC degrader, in metastatic castration-resistant prostate cancer (mCRPC). American Society of Clinical Oncology: 2022.

(10) Chamberlain, P. P.; D”Agostino, L. A.; Ellis, J. M.; Hansen, J. D.; Matyskiela, M. E.; McDonald, J. J.; Riggs, J. R.; Hamann, L. G. Evolution of Cereblon-Mediated Protein Degradation as a Therapeutic Modality. ACS Med Chem Lett 2019, 10 (12), 1592–1602. DOI: 10.1021/acsmedchemlett.9b00425 From NLM PubMed-not-MEDLINE.

(11) Schneekloth, A. R.; Pucheault, M.; Tae, H. S.; Crews, C. M. Targeted intracellular protein degradation induced by a small molecule: En route to chemical proteomics. Bioorg Med Chem Lett 2008, 18 (22), 5904–5908. DOI: 10.1016/j.bmcl.2008.07.114 From NLM Medline.

(12) Burslem, G. M.; Smith, B. E.; Lai, A. C.; Jaime-Figueroa, S.; McQuaid, D. C.; Bondeson, D. P.; Toure, M.; Dong, H.; Qian, Y.; Wang, J.; et al. The Advantages of Targeted Protein Degradation Over Inhibition: An RTK Case Study. Cell Chem Biol 2018, 25 (1), 67–77 e63. DOI: 10.1016/j.chembiol.2017.09.009 From NLM Medline.

(13) Winter, G. E.; Buckley, D. L.; Paulk, J.; Roberts, J. M.; Souza, A.; Dhe-Paganon, S.; Bradner, J. E. DRUG DEVELOPMENT. Phthalimide conjugation as a strategy for in vivo target protein degradation. Science 2015, 348 (6241), 1376–1381. DOI: 10.1126/science.aab1433 From NLM Medline.

(14) Burslem, G. M.; Song, J.; Chen, X.; Hines, J.; Crews, C. M. Enhancing Antiproliferative Activity and Selectivity of a FLT-3 Inhibitor by Proteolysis Targeting Chimera Conversion. J Am Chem Soc 2018, 140 (48), 16428–16432. DOI: 10.1021/jacs.8b10320 From NLM Medline.

(15) Liu, Z.; Hu, M.; Yang, Y.; Du, C.; Zhou, H.; Liu, C.; Chen, Y.; Fan, L.; Ma, H.; Gong, Y.; Xie, Y. An overview of PROTACs: a promising drug discovery paradigm. Mol Biomed 2022, 3 (1), 46. DOI: 10.1186/s43556-022-00112-0 From NLM PubMed-not-MEDLINE.

(16) Bricelj, A.; Steinebach, C.; Kuchta, R.; Gutschow, M.; Sosic, I. E3 Ligase Ligands in Successful PROTACs: An Overview of Syntheses and Linker Attachment Points. Front Chem 2021, 9, 707317. DOI: 10.3389/fchem.2021.707317 From NLM PubMed-not-MEDLINE.

(17) Bemis, T. A.; La Clair, J. J.; Burkart, M. D. Unraveling the Role of Linker Design in Proteolysis Targeting Chimeras. J Med Chem 2021, 64 (12), 8042–8052. DOI: 10.1021/acs.jmedchem.1c00482 From NLM Medline.

(18) Cyrus, K.; Wehenkel, M.; Choi, E. Y.; Han, H. J.; Lee, H.; Swanson, H.; Kim, K. B. Impact of linker length on the activity of PROTACs. Mol Biosyst 2011, 7 (2), 359–364. DOI: 10.1039/c0mb00074d From NLM Medline.

(19) Poongavanam, V.; Kolling, F.; Giese, A.; Goller, A. H.; Lehmann, L.; Meibom, D.; Kihlberg, J. Predictive Modeling of PROTAC Cell Permeability with Machine Learning. ACS Omega 2023, 8 (6), 5901–5916. DOI: 10.1021/acsomega.2c07717 From NLM PubMed-not-MEDLINE.

(20) Matsson, P.; Doak, B. C.; Over, B.; Kihlberg, J. Cell permeability beyond the rule of 5. Adv Drug Deliv Rev 2016, 101, 42–61. DOI: 10.1016/j.addr.2016.03.013 From NLM Medline.

(21) Cantrill, C.; Chaturvedi, P.; Rynn, C.; Petrig Schaffland, J.; Walter, I.; Wittwer, M. B. Fundamental aspects of DMPK optimization of targeted protein degraders. Drug Discov Today 2020, 25 (6), 969–982. DOI: 10.1016/j.drudis.2020.03.012 From NLM Medline.

(22) Edmondson, S. D.; Yang, B.; Fallan, C. Proteolysis targeting chimeras (PROTACs) in “beyond rule-of-five” chemical space: Recent progress and future challenges. Bioorg Med Chem Lett 2019, 29 (13), 1555–1564. DOI: 10.1016/j.bmcl.2019.04.030 From NLM Medline.

(23) Apprato, G.; Ermondi, G.; Caron, G. The Quest for Oral PROTAC drugs: Evaluating the Weaknesses of the Screening Pipeline. ACS Med Chem Lett 2023, 14 (7), 879–883. DOI: 10.1021/acsmedchemlett.3c00231 From NLM PubMed-not-MEDLINE.

(24) Doak, B. C.; Zheng, J.; Dobritzsch, D.; Kihlberg, J. How Beyond Rule of 5 Drugs and Clinical Candidates Bind to Their Targets. J Med Chem 2016, 59 (6), 2312–2327. DOI: 10.1021/acs.jmedchem.5b01286 From NLM Medline.

(25) Doak, B. C.; Over, B.; Giordanetto, F.; Kihlberg, J. Oral druggable space beyond the rule of 5: insights from drugs and clinical candidates. Chem Biol 2014, 21 (9), 1115–1142. DOI: 10.1016/j.chembiol.2014.08.013 From NLM Medline.

(26) Veber, D. F.; Johnson, S. R.; Cheng, H. Y.; Smith, B. R.; Ward, K. W.; Kopple, K. D. Molecular properties that influence the oral bioavailability of drug candidates. J Med Chem 2002, 45 (12), 2615–2623. DOI: 10.1021/jm020017n From NLM Medline.

(27) DeGoey, D. A.; Chen, H. J.; Cox, P. B.; Wendt, M. D. Beyond the Rule of 5: Lessons Learned from AbbVie”s Drugs and Compound Collection. J Med Chem 2018, 61 (7), 2636–2651. DOI: 10.1021/acs.jmedchem.7b00717 From NLM Medline.

(28) Price, E.; Weinheimer, M.; Rivkin, A.; Jenkins, G.; Nijsen, M.; Cox, P. B.; DeGoey, D. Beyond Rule of Five and PROTACs in Modern Drug Discovery: Polarity Reducers, Chameleonicity, and the Evolving Physicochemical Landscape. J Med Chem 2024. DOI: 10.1021/acs.jmedchem.3c02332 From NLM Publisher.

(29) Ermondi, G.; Vallaro, M.; Goetz, G.; Shalaeva, M.; Caron, G. Updating the portfolio of physicochemical descriptors related to permeability in the beyond the rule of 5 chemical space. Eur J Pharm Sci 2020, 146, 105274. DOI: 10.1016/j.ejps.2020.105274 From NLM Medline.

(30) Weiss, D. R.; Baylon, J. L.; Evans, E. D.; Paiva, A.; Everlof, G.; Cutrone, J.; Broccatelli, F. Balanced Permeability Index: a multi-parameter index for improved in-vitro permeability. bioRxiv 2023, 2023.2012.2001.569478. DOI: 10.1101/2023.12.01.569478.

(31) Pond, S. M.; Tozer, T. N. First-Pass Elimination Basic Concepts and Clinical Consequences. Clinical Pharmacokinetics 1984, 9 (1), 1–25. DOI: 10.2165/00003088-198409010-00001.

(32) Burton, P. S.; Goodwin, J. T.; Vidmar, T. J.; Amore, B. M. Predicting drug absorption: how nature made it a difficult problem. J Pharmacol Exp Ther 2002, 303 (3), 889–895. DOI: 10.1124/jpet.102.035006 From NLM Medline.

(33) Tanaka, Y.; Kitamura, Y.; Maeda, K.; Sugiyama, Y. Explication of Definitional Description and Empirical Use of Fraction of Orally Administered Drugs Absorbed From the Intestine (Fa) and Intestinal Availability (Fg): Effect of P-glycoprotein and CYP3A on Fa and Fg. J Pharm Sci 2016, 105 (2), 431–442. DOI: 10.1016/j.xphs.2015.11.005 From NLM Medline.

(34) Zhao, Y.; Jona, J.; Chow, D. T.; Rong, H.; Semin, D.; Xia, X.; Zanon, R.; Spancake, C.; Maliski, E. High-throughput logP measurement using parallel liquid chromatography/ultraviolet/mass spectrometry and sample-pooling. Rapid Commun Mass Spectrom 2002, 16 (16), 1548–1555. DOI: 10.1002/rcm.749 From NLM Medline.

(35) Goetz, G. H.; Farrell, W.; Shalaeva, M.; Sciabola, S.; Anderson, D.; Yan, J.; Philippe, L.; Shapiro, M. J. High throughput method for the indirect detection of intramolecular hydrogen bonding. J Med Chem 2014, 57 (7), 2920–2929. DOI: 10.1021/jm401859b From NLM Medline.

(36) Bower, M. J.; Aronov, A. M.; Cleveland, T.; Hariparsad, N.; McGaughey, G. B.; McMasters, D. R.; Zhang, X.; Goldman, B. Smallest Maximum Intramolecular Distance: A Novel Method to Mitigate Pregnane Xenobiotic Receptor Activation. J Chem Inf Model 2020, 60 (4), 2091–2099. DOI: 10.1021/acs.jcim.9b00692 From NLM Medline.

