## Supporting Information for "Balanced Permeability Index is a Strong Predictor of Intestinal Absorption and Oral Bioavailability for Heterobifunctional Ligand-Directed Degraders"


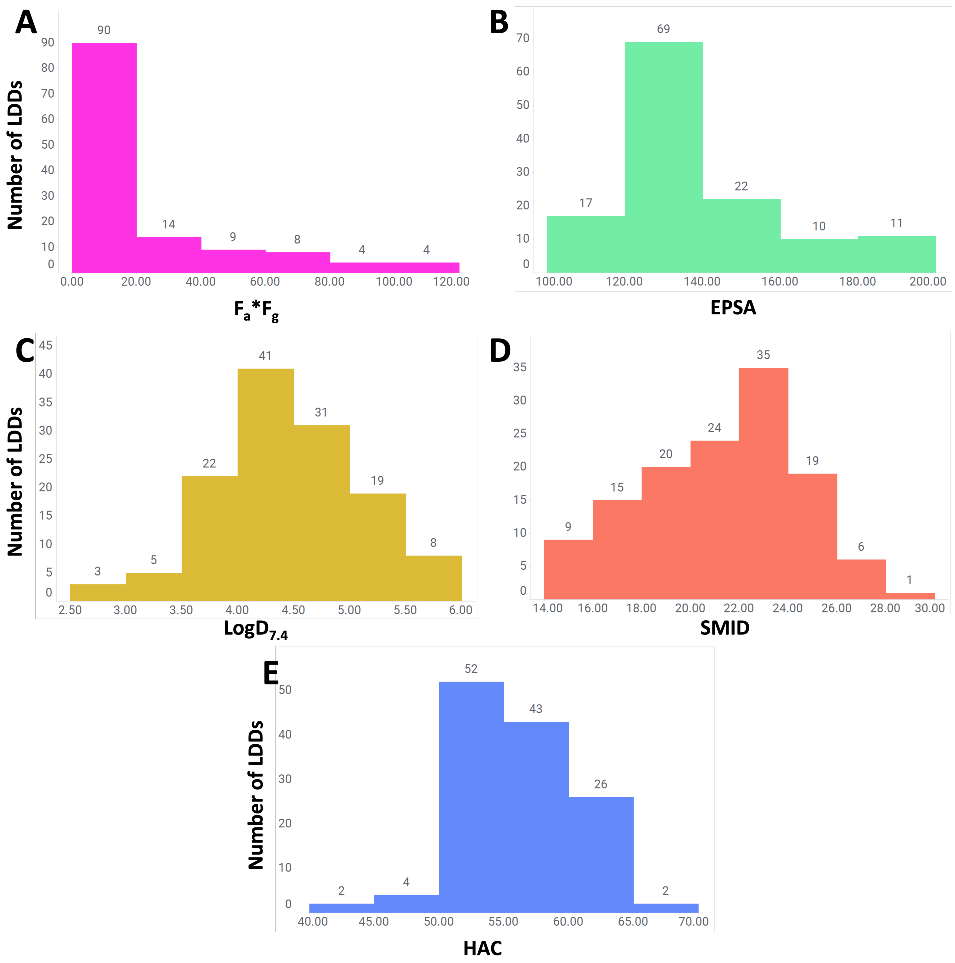


Figure S1 Distributions of (A) $F_a*F_g$ and key $BPP_{LDD}$ components in out 129 LDD internal dataset.

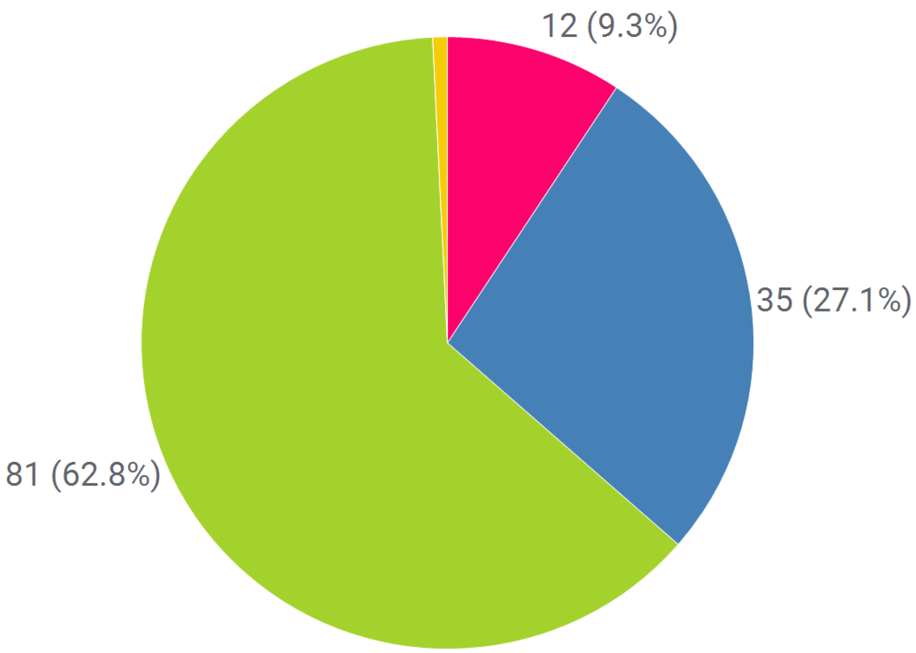


Figure S2. Breakdown of how many compounds per program are in our 129 LDDs internal $F_{a*}F_g$ dataset.

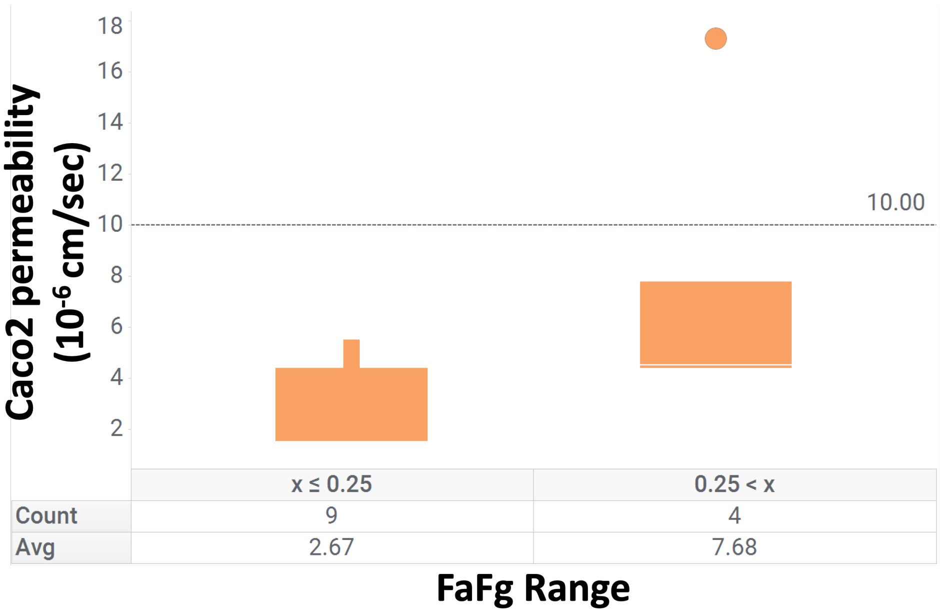


Figure S3. Distribution of Caco-2 Papp A-to-B binned by $F_a{*}F_g$ for LDDs with recovery from the Caco-2 assay > 80% in our dataset.

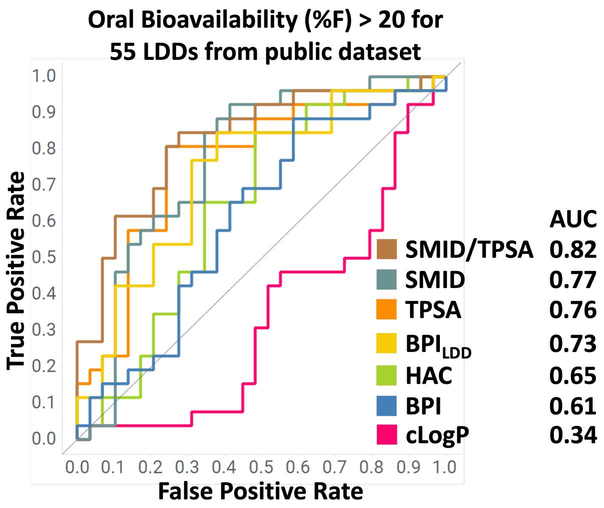


Figure S4. ROC analysis using a public dataset of 55 LDDs with oral bioavailability.[1] For this set the high %F cutoff was set at 20 following a previous study.[2] AUC value for each parameter is presented in the legend.

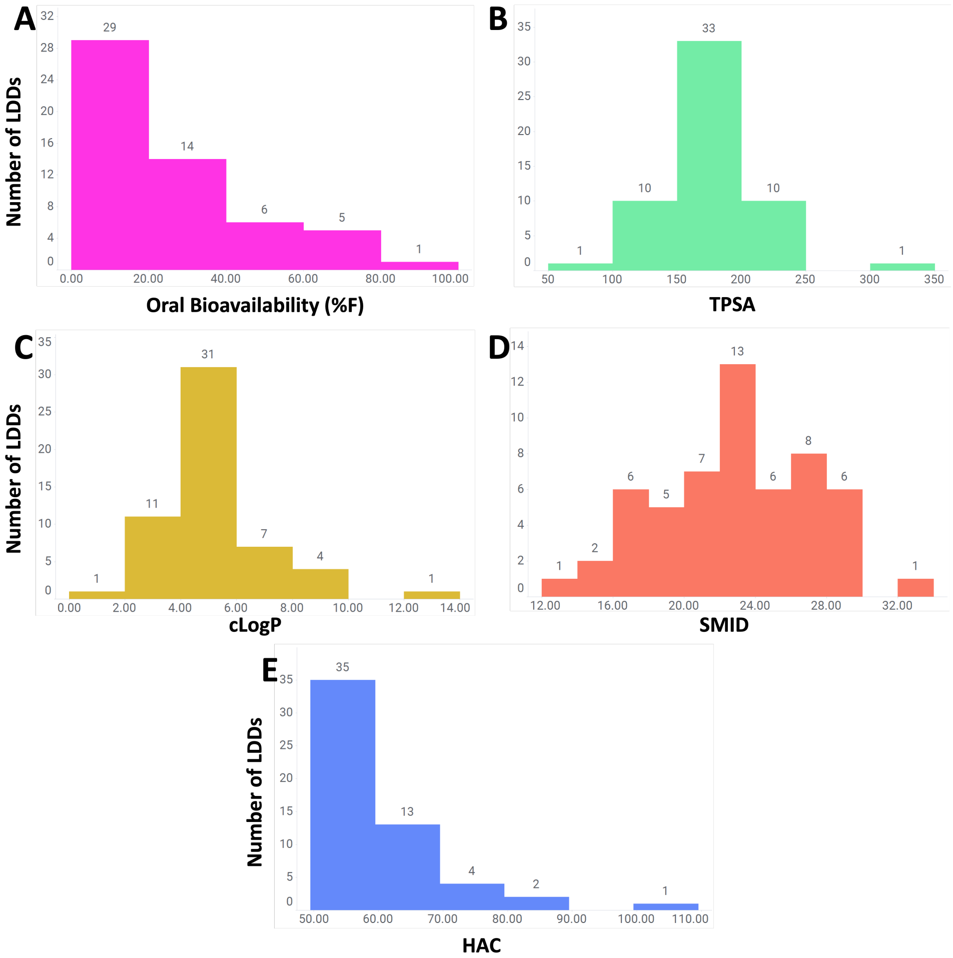


Figure S5. Distribution of (A) oral bioavailability (%F) and key BPILDD paremeters for the 55 LDD public dataset.
